# Chrombus-XMBD: A Graph Generative Model Predicting 3D-Genome, *ab initio* from Chromatin Features

**DOI:** 10.1101/2023.08.02.551072

**Authors:** Yuanyuan Zeng, Zhiyu You, Jiayang Guo, Jialin Zhao, Ying Zhou, Jialiang Huang, Xiaowen Lyu, Longbiao Chen, Qiyuan Li

## Abstract

The landscape of 3D-genome is crucial for transcription regulation. But capturing the dynamics of chromatin conformation is costly and technically challenging. Here we described “Chrombus-XMBD”, a graph generative model capable of predicting chromatin interactions *ab inito* based on available chromatin features. Chrombus employes dynamic edge convolution with QKV attention setup, which maps the relevant chromatin features to a learnable embedding space thereby generate genomewide 3D-contactmap. We validated Chrombus predictions with published databases of topological associated domains (TAD), eQTLs and gene-enhancer interactions. Chrombus outperforms existing algorithms in efficiently predicting long-range chromatin interactions. Chrombus also exhibits strong generalizability across different cell lineage and species. Additionally, the parameter sets of Chrombus inform the biological processes underlying 3D-genome. Our model provides a new perspective towards interpretable AI-modeling of the dynamics of chromatin interactions and better understanding of *cis*-regulation of gene expression.

## Introduction

The three-dimensional conformation of chromatin, or 3D-genome, are crucial for organization and regulation of gene transcription[1]. Formation of DNA looping structures mediated by specific epigenetic modifications and trans-acting proteins is the molecular basis for various *cis*-regulation activities in eukaryote cells, such as enhancer-promoter and enhancer-enhancer interactions[2, 3]. Techniques such as 3C, 4C and Hi-C are developed to investigate DNA interactions at either specific locus or at a genome-wide level. These interactions can be used to interpret the biological processes underlying known trait associated loci (TAL), such as GWAS risk loci and eQTL, which are widely implicated in development, differentiation and diseases[4-7]. The most common structural elements of 3D-genome are DNA loops. DNA loops enable interaction between two separated DNA fragments, which is required by many *cis*-regulatory elements[8, 9]. The formation of DNA loops is a complex process which involves not only covalent bonding of DNA but also locus-specific binding of proteins[10, 11]. According to the loop extrusion hypothesis, Cohesin, a protein complex encircling DNA, actively slides along the chromatin fiber and extrude the latter into a circular conformation[12, 13]. The extrusion process halts when Cohesin encounters with CTCF at convergent CTCF-binding sites, resulting in stabilized DNA loops[14, 15]. The stationary distribution of hundreds of thousands of DNA loops form a specific landscape, in which chromatin is segregated into many topological associated domains (TAD) of highly frequent DNA interactions[16]. TADs are further aggregated into large chromosomal compartments. A Compartments are linked to active chromatin and gene expression, while B compartments are associated with repressed chromatin and gene silencing[17, 18].

To date, our understanding of the 3D-genome is primarily based on Hi-C experiments conducted in cell lines or tissues. Many computational methods are derived to infer 3D-genome which yielded insights into the *cis*-regulatory program of eukaryote cells. Nevertheless, most of the studies are still based on limited sample size, and the algorithms often lack consistency, which hinders the understanding of 3D-genome organization under more diverse biological contexts. Recent studies use machine learning methods to predict 3D-genome from chromatin features, which provide an alternative approach to solve the complex chromatin conformation. Akita [19] and DeepC [20] used deep convolutional neural network to predict locus-specific genome folding from DNA sequence alone. HiC-Reg[21] and Epiphany[22] predict Hi-C-based contact counts using chromatin features, such as DNase I hypersensitive sites, CTCF, H3K27ac, H3K27me3 and H3K4me3 at a resolution of 5kb. These methods have yielded decent prediction accuracy for 3D chromatin organization, with Pearson correlation coefficients ranging from 0.57 to 0.76.

Yet most of these methods consider the chromatin features of each evenly binned segment are independent to each other. In reality, however, chromatin segmentation is no evenly distributed along the genome, and the interactions between two DNA segments can be influenced by chromatin features of their distant neighbors[8, 23]. Although increase of the bin size can capture more information of the landscape of certain features, it also downscales the resolution and expands the parameter space. Nevertheless, the predictions of these methods are all limited to 1Mb, hence cannot predict long-range DNA interactions which are often more important in biology.

Many complex biological interactions are represented by graphs[24-26]. Graph neural network (GNN) provides a framework for knowledge inference on graphs, which is successfully applied to drug discovery, protein conformation and gene expression analysis[27, 28]. The advantages of GNN in modeling biological systems rely on its capability to process irregular, complex interactions among biological entities.

Here we described a new, graph-based generative model, “Chrombus”, which is capable for *ab initio* prediction of the 3D conformation of chromatin based on specific epigenetic features. Our algorithm aggregates signals from a set of neighboring CTCF segments and predicts the interaction potential thereby reveal context-specific formation of TAD. Of note, our graph-based algorithm showed highly generalizable predictive power and capable of predicting long-range DNA interactions, which outperform the state-of-the-art methods [19-22].

We rigorously validated our method against known evidences of 3D-genome, including eQTL associations, enhancer-promoter architectures. We demonstrated that “Chrombus” can recover known *cis*-regulatory events and predict regulatory programs highly relevant to biology. Our method provides an efficient, robust way to reconstruct the landscape of 3D-genome at all levels.

## Data and Methods

### Datasets

The training and validation of “chrombus” are based on data of three cell lines, GM12878, K562 and H1ESC (Supplementary Table 1). GM12878 is used for model training and testing. K562 and CH12 were used for external validation and evaluation of model generalization.

For each cell line, six epigenetic features were collected from the Encyclopedias of DNA Elements (ENCODE)[29] and 4D Nucleome (4DN) projects[30] including CTCF, RAD21 (cohesin), H3K27ac, H3K4me3, POLR2A, open chromatin (DNase I). All data were based on published ChIP-Seq analysis, preprocessed and QC’ed follow the standard processes of data consortia. Preprocessed Hi-C data for GM12878, K562 and CH12 were obtained from GSE63525[31]. All Hi-C data are scaled to the same resolution of 5000bp. All chromosomal coordinates are based on hg19 reference genome build.

### Graph representation of 3D-genome based on CTCF-segments

We segmented the chromatin into uneven fragments based on CTCF-binding peaks. Each fragment corresponds to a vertex, denoted as ***V***_***i***_ ∈ ***V*** (Supplementary Table 1). Then the edge, ***E***_***ij***_, between the vertices *i* and *j* were defined by the averaged interaction score between the two fragments derived directly from processed Hi-C data. Thus, the 3D-structure of the chromatin is represented by a graph ***G*(*V, E*)** (Figure 1). Each vertex is represented by 14-dimension feature vector ***x***_***i***_. The features include CTCF-binding strength at both ends of the segment, denotes as left peak and right peak. Then we inferred the directionality of CTCF motifs within the CTCF-binding sites using fimo[32]. Each CTCF-site was labeled as “0”, “1” according to the positive and negative strand. The RAD21-binding at a given CTCF-site (“left cohesin” and “right cohesin”) was also processed as a binary status, where “1” indicated that RAD21-binding peak was in vicinity of the CTCF-site (within a 500bp). Furthermore, we incorporated mean value of H3K27ac, H3K4me3, POLR2A and DNase I signals to represent the epigenetic character of each segment, which were scaled to the range of [0, 1]. Finally, we included the chromosomal coordinates of both ends of a segment, resulting in a total of 14 node attributes (Supplementary Table 1, Supplementary Figure2A-C). As the edges in the graph depends on the strength of interaction, we filtered out edges of weak interaction (raw Hi-C value > 0). To define the edge attributes, we calculated the mean contact value of all 5kb-bin located in each segment. There are three methods for normalizing the contact values: SQRTVC, KR and VC. To determine the most appropriate normalization method, we chose the normalization method which best discriminate interactions within the same TAD (“within-TAD”) and those spanning different TADs (“between-TAD”) by cross-entropy. A higher cross-entropy indicates a better discriminative power between the two groups. The TADs were inferred from normalized Hi-C data using the Arrowhead algorithm[31]. Based on our evaluation, SQRTVC was chosen as normalization method (Supplementary Figure 1).

**Figure 1.**
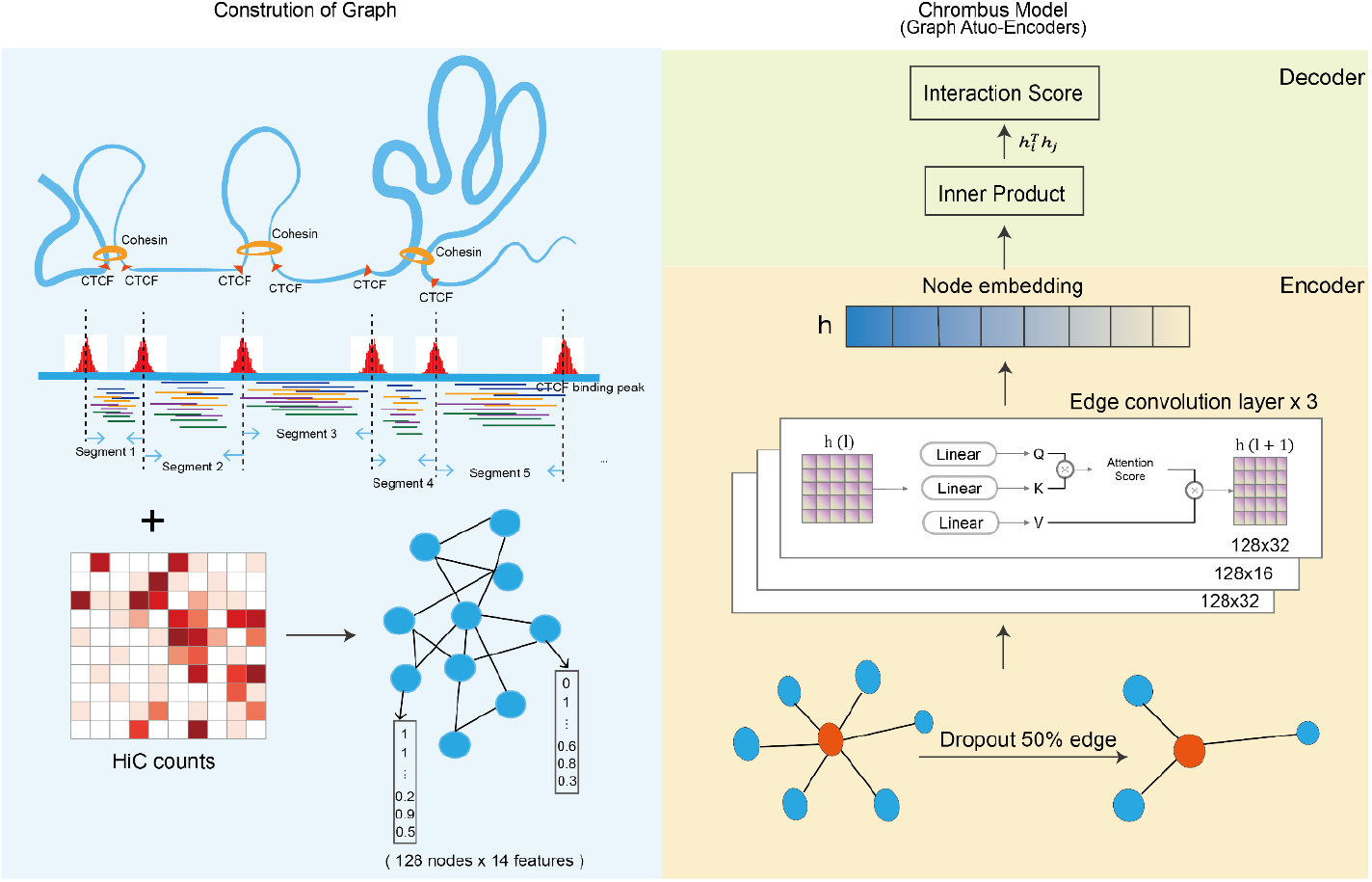
Construction of graph and model architecture. Each graph consists 128 vertices, and each vertex represents a chromatin segment derived from CTCF binding peaks. The node (vertex) attributes consist 14-dimensional chromatin features. The goal of the learning process is to generate the interactions among vertices, of which the labelling is based on Hi-C data (left panel). Chrombus is adopted from GAE architecture. The encoder consists of three edge convolution layers with embedded QKV attention mechanism and outputs embedding of dimensions 32, 16, and 32. The decoder is implemented as a plain inner product (right panel).

### The Chrombus Model

We developed “Chrombus”, a generative graph convolution model which reconstructs chromatin interaction (Figure 1). Chrombus adopts a typical GAE[33] model. The encoder maps 14-dimensional features of a graph of *N* vertices ***x*** = (*x*_1_, …, *x*_*N*_) to 32-dimensional representation ***z*** = (*z*_*i*_, …, *z*_*N*_) through 3-depth dynamic edge-convolution layers. And layer 3 is stepped over a single nonlinear layer from to layer 1. Given ***z***, the decoder then generates *N*-by-*N* matrix of interaction strength as ***A*** = ***z′z***, for each pair of vertices.

The graph convolution layers follow an architecture of stacked self-attention, dynamic edge convolution and fully connected layers for encoder, and dot product for decoder (Figure 1).

The encoder takes batched subgraphs as input. Each subgraph contains 128 vertices, representing 128 consecutive CTCF-segments which were randomly cropped from a chromatin. During training and testing, the vertices were connected by random edges based on following rules. 1) no interactions beyond the maximal span of 32 consecutive CTCF-segments; and 2) within the maximal span, any pair of CTCF-segments are connected by 50% chance, which correspond to an Erdős-Rényi random graph of G(128, 0.5).

The encoder consists three edge-convolution layers, each of which transform the node features from the previous layer by incorporating messages from random assigned neighbors. The output dimensions of each layer are 32, 16, 32, respectively (Figure 1).

Let normalized features of the target node *i* in the first layer 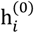 be the transformed from input feature 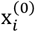 and features of the neighboring source nodes *j* ∈ N(*i*) (eq. 1).

Subsequently, the hidden variable of the *l* -th layer, 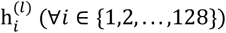 is obtained by *l*-th (∀*l* ∈ {1,2,3}) graph convolution operator:

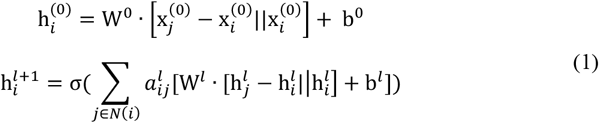

Where W^*l*^ and b^*l*^ denotes the trainable weights and bias of the *l* -th layer, σ represents the sigmoid activation function, *N*(*i*) represents the neighbors of node *i, i* and *j* represent the indices of target and source nodes, respectively. 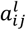 is partition matrix that gathers and reallocates the weight of hidden representations from *N*(*i*) in the *l*-th layer.

To ensure the target node receives relevant information from its random neighbors during edge-convolution, we employed a multi-head (n=8) self-attention as described in transformer model[34] (eq. 2). We used three 1-depth linear layers to determine query, key and value. The attention score 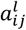 is defined as follow:

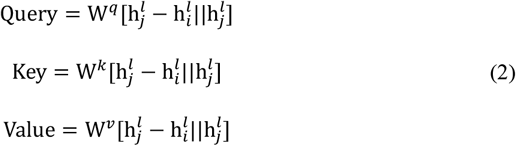

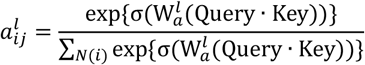

Where 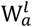 represents the weights of attention, and [· || ·] denotes the concatenation of multiple vectors. The increment of hidden expression of node *j* over its center node *i* is passed through the edge when calculating 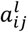.

In addition to self-attention, we design a signed edge weight to address the biological fact that interaction between segments is negatively correlated with the linear distance (eq. 3). We defined a *cis*-interaction range of 9 consecutive segments so that the sign of *ω* flips according to *D*_*ij*_, the linear distance between two segments measured by number of segments in between.

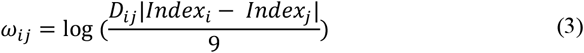

### Decoder

To acquire the interaction scores among nodes, we apply inner production to the aforementioned embedding of each pair of all nodes, which is as follow:

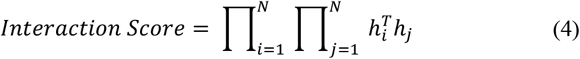

Where the *Interaction Score* represents the interaction strength inferred by decoder between node i and node j. The mean square error (MSE) is broadly used for regression problems. Eventually, our objective of model is to minimize MSE between predicted and real interaction of target nodes.

### Model training and testing

Each sample for GM12878 model is randomly put-back cropped into 128-segments from the training and testing chromosomes. For model training, 200 samples were generated from each chromosome, and 50 samples for testing purposes.

Regarding the generalization model, chromosomes are cropped into non-overlapping samples using a 128-segment window. In total, 301 (GM12878), 420 (K562) and 455 (CH12) samples are generated. Out of these samples, 50 samples are randomly selected for testing. To train the model, a five-fold cross-validation approach is applied, utilizing the least number of samples. Four model are trained, and the model with the best performance on the testing set is selected for cross-cell prediction.

### Comparing interactions located Within TAD and between TAD

TADs were assessed using Arrowhead and HiCexplorer[35]. Interactions between two segments within a TAD were categorized as “Within TAD”, while the interactions between two segments located in different adjacent TADs were categorized as “Between TAD”. For TAD defined by both methods, we examined the chromatin activities of these two groups according to the Chrombus’s predictions across 22 autosomes. Receiver operating characteristic (ROC) curves were generated, and the area under the curve (AUC) values were calculated to assess the ability of interaction scores to distinguishing between the two groups. The AUC value and confidence interval for the HiC scores and Predicted scores were calculated using the ROCit R package. We performed an Inferior test to compare the predicted score to HiC scores using rocNIT R package.

### Enrichment of eQTLs, enhancer-gene interactions

eQTLs of GM12878 with hg19 coordinates were download from GTEX (Version 7)[36]. Enhancer-gene interactions were obtained from GeneHancer[37]. The eQTLs and enhancer-gene interactions are referred as interaction events. We categorized each pair of segments into two groups: those encompass interaction events and those do not. We then compared the distribution of chromatin interaction scores between these two groups, and the significance of the difference was evaluated using the Wilcoxon test.

To calculated the fold of enrichment, we considered the total number of predicted segment pairs as *N* and the total number of segment pairs encompassing interactions as *n*. The background ratio was calculated as *n*/*N*. Then we applied incremental thresholding to the predicted scores, for all segments with predicted scores above the threshold value, we calculated the ratio of segment pairs encompassing interaction events (*n*^′^/*N*^′^). The enrichment fold was then calculated by (*n*^′^/*N*^′^)/(*n*/*N*).

### Comparing Chrombus with HiC-Reg

We obtained cross-chromosome predictions for GM12878 from HiC-Reg, specifically for chromosome 14 and 17 [21]. To process the data, we converted the contact counts of 5-kb bins into segment-based contact counts by taking the mean value. The interactions were then divided into three groups based on their distance: within 0.5M, between 0.5-1M and beyond 1M.

Within each distance group, we calculated the Pearson correlation coefficient between the HiC scores and predicted scores from HiC-Reg and Chrombus. All the calculations and visualizations were performed using R (version 4.1.2).

### Feature importance and node representation

The input features’ contributions in predicting chromatin interaction are computed via the GNNexplainer algorithm. The node mask value is learned using PyTorch genometric (v2.3.1) with 200 epochs. The correlation between node representation and input feature is determined using the Pearson correlation coefficient.

## Results

### Chrombus: Accurate prediction of chromatin interactions using epigenetic data through a graph neural network

We developed “Chrombus”, a graph-based generative model to predict chromatin interactions *ab initio* based on epigenetic features. “Chrombus” used three dynamic edge-convolution layers with multi-head attention, and was trained with multi-level epigenetic data (as input) and Hi-C data (as label) of lymphoblastoid (GM12878) Hi-C. Instead of evenly binning the chromatin, we used CTCF binding peaks to define segments of DNA as units of interactions. A typical dataset contains 39,998-59,443 segments (Supplementary Figure 2D).

The training data consist batched subgraphs of 128 adjacent CTCF-segments which were randomly cropped from the contact map (Methods). In each round of training, one of 22 autosomes was left out as test data with the rest as training data. To evaluate the predict performance, the training-test process was repeated 22 rounds until each autosome was tested independently based on model trained on the others, which resulted in 22 models.

Each model reached convergence after approximately 400 epochs. Both the training losses and the testing losses followed the same trend during the training process (Supplementary Figure 3A). We evaluated the performance of “Chrombus” using Pearson’s correlation coefficients between the predicted scores and the actual HiC scores. For the training data, all 22 models converged with correlation coefficients ranging from 0.880 to 0.893 and the mean square error (MSE) ranging from 0.140 to 0.162. As For the test data, the correlation coefficients ranging from 0.849 to 0.900 and the mean square error (MSE) ranging from 0.126 to 0.286 (Figure 2A). We also noticed that the training correlation and testing correlation decrease slightly with the sizes of the CTCF-segments (Figure 2B).

**Figure 2.**
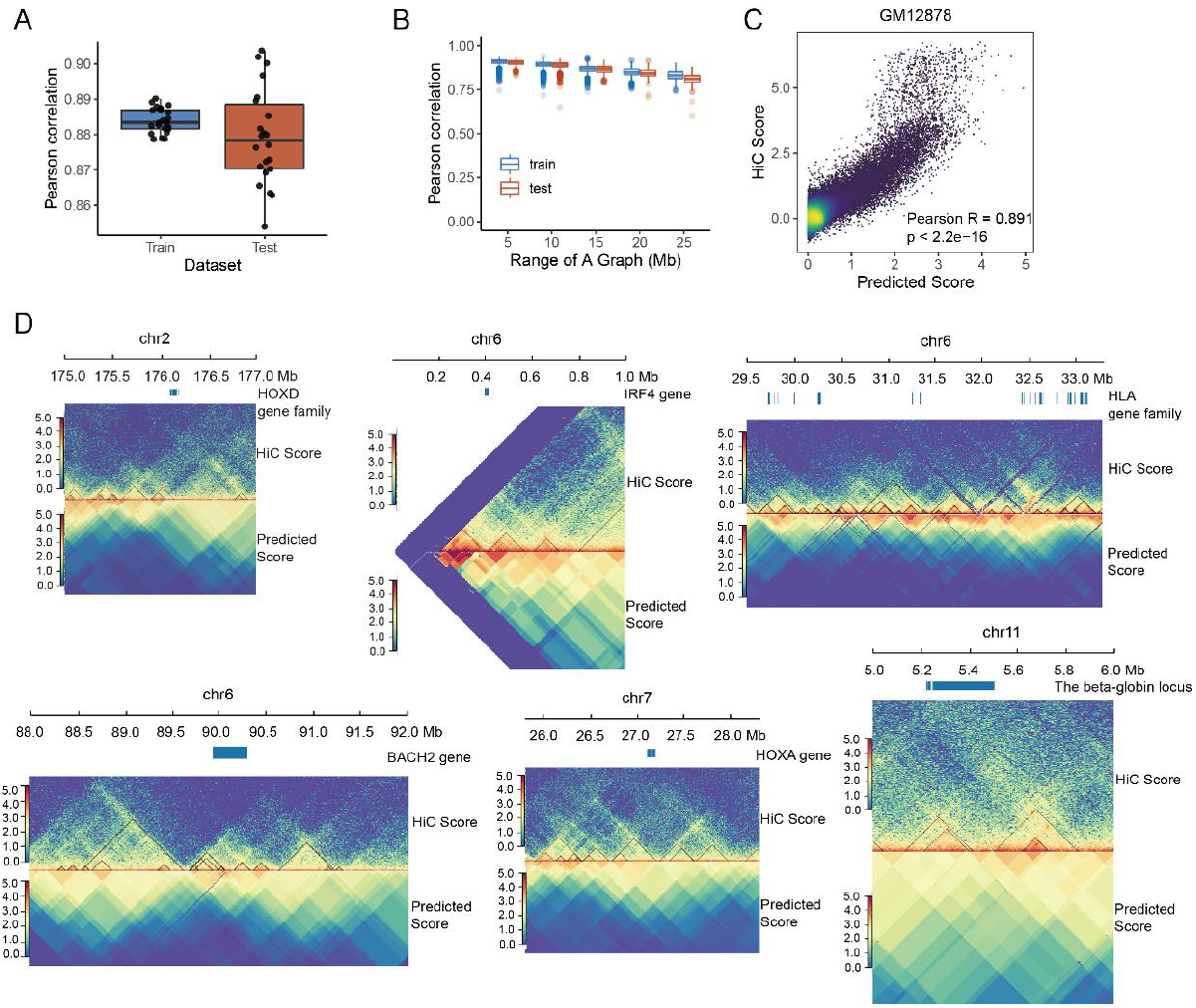
Chrombus recover known chromatin interactions and TAD. The model parameters were trained and tested independently using chromatin features of GM12878. The consistency between the true Hi-C values and the predicted values was evaluated using Pearson’s correlation coefficient. A: Distribution of correlation coefficients between Hi-C values and the predicted values of Chrombus in the training and test datasets. B: The distribution of correlation coefficients for chromatin interactions of different distances. C: Scatter plot of randomly selected 100,000 predicted values against true Hi-C values. D: predictions at six known TAD loci.

We further estimated the overall correlation between predicted scores and HiC scores in 100,000 (2.169%) randomly sampled pairs from all 22 autosomes and yielded a correlation coefficient of 0.891 (0.889-0.892, 95% CI) (Figure 2C).

To cope with the distance and size effects of 3D genome, we devised a contrast edge weight (9) based on average number segment counts within TAD. When two segments are separated by more than 9 other segments, self-attention assign positive weights to stronger interactions. Conversely, when two segments are located within 9 segments, self-attention assign negative weights to stronger interactions. This approach enhances the model’s sensitivity to long-distance interactions, which hold greater biological significance, and dissociation events that occur between segments of shorter distances, commonly found at the boundaries of TAD or compartments. From the testing results, we selectively examine a set of genes related to chromatin 3D structures in GM12878 cell line, including the TADs located at IRF4[38], BACH [39], HOXD family[40], HLA family[41], HOXA family[42] and the beta-globin locus[43] (Figure 2 D). These TADs spanned various regions of genome, and the predictions of Chrombus highly correlated to the corresponding Hi-C scores, with Pearson’s correlation coefficients ranging from 0.71 to 0.94 (Supplementary Figure 3B). The results suggest Chrombus is capbale to recover known TADs with high consistency.

### Chrombus recovers DNA-DNA interaction landscape as represented by TAD

Topologically associating domains (TAD) are known as the elementary three-dimensional structure of chromatin with a crucial role in transcriptional regulation. TADs are isolated from each other at the boundary ends. More active DNA-DNA interactions occur within the TAD than between, which resulting in topology associated transcription regulation of genes. Leveraging this distinctive feature, we further verified Chrombus predictions within and between TADs.

We opted for two widely used methods, “HiCexplorer” and “Arrowhead”, to define TADs from HiC data (Methods). Consequently, we categorized all predicted chromatin interactions into two classes, namely, “within TAD” and “between TAD”. Based on the reference TAD derived by ArrowHead, the AUC of HiC scores and Chrombus predicted scores are 0.928 and 0.832, respectively (Figure 3A). And for HiCexplorer -derived reference TADs, the AUC values are 0.861 and 0.766 for HiC and Chrombus (Figure 3B). In both cases, Chrombus predictions are capable to distinguish within and between TAD with non-inferiority (P < 0.05).

**Figure 3.**
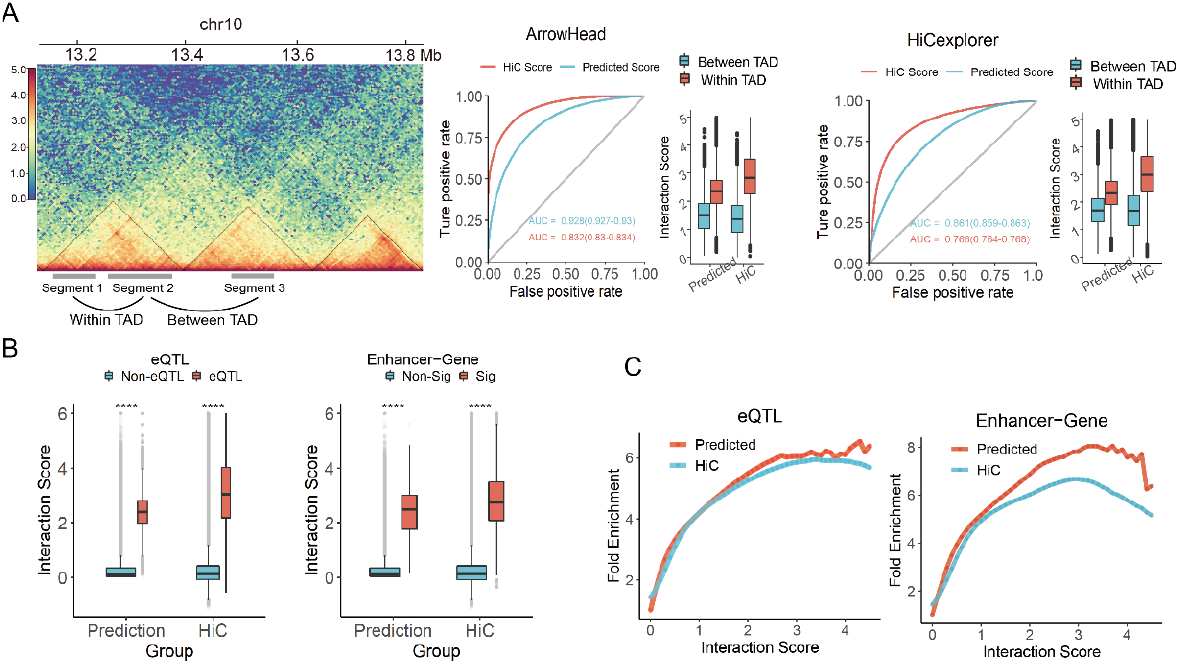
Validation of Chrombus predictions by published datasets of TADs, eQTLs and enhancer-gene interactions. A: illustration of interactions within and between TAD (left). ROC curves for classifying chromatin interactions within and between TAD based on predicted scores and Hi-C scores, along with interaction strength in both groups. As reference, TAD in GM12878 were inferred from the same Hi-C data using Arrowhead and HiCexplorer. B: Predicted and true Hi-C interactions of segment-pairs encompassing eQTL (left) and significant enhancer-gene interactions (right) against those do not. C: Fold of enrichment of eQTL and enhancer-gene interactions in segment pairs of various thresholding of either predicted or Hi-C based interaction scores.

3D chromatin enables interactions among different *cis*-elements, which underlie the function of expression quantitative trait loci (eQTL). Hence we validated Chrombus predictions against known eSNP-eGENE interactions in normal tissues[36] and enhancer-gene interactions from the published database[37] For comparison, we also evaluated the association of HiC scores to these interactions. As a result, in pairs of CTCF-segments encompassed eSNP-eGENE interactions and enhancer-gene interactions, the predicted scores by Chrombus are significantly higher than the background (P < 0.05, Figure 3B). Similar tendency can be observed in HiC signals. Furthermore, we found that the enrichment of both types of interactions increases with the Chrombus predicted scores as well as the HiC signals (Figure 3C). Notably, as the scores increase, the fold-of-enrichment of the interactions based on Chrombus prediction surpassed that based on the HiC signals, suggesting Chrombus is capable to better recover the *cis*-regulatory activities than original HiC.

Thus, our study not only elucidates the association between chromatin 3D structure and distant interactions events, but also demonstrates the agreement between model predictions and HiC scores from a fine-grained perspective.

### Chrombus is generalizable cross cell lineage and species

While other models require training and prediction in the same cell line, Chrombus is highly generalizable cross different cell types and lineage. Here, we examine Chrombus’s generalizability on GM12878, K562 and CH12 cell lines. The model of each cell line that exhibited the best performance in the cross-validation were selected for cross-cell line and cross-species prediction (supplementary Figure 4).

In the three cell lines, the models of the best performance in cross-validation reached Pearson’s correlation coefficients of 0.71 (CH12), 0.82 (GM12878), and 0.80 (K562) between HiC signals and the predicted scores. When we applied the models trained on one cell line to the other two, we observed trivial decrease in the performance between two human cell lines (GM12878 and K562), but significant decrease in the performance in mouse-derived cell line (CH12) (Figure 4A). For instance, the model trained on GM12878 reached a correlation of 0.8 in K562, whereas the correlation in CH12 is just 0.59. In contrast, the mouse model showed relatively stable performance in two human cell lines. These findings suggest that Chrombus are capable to capture conserved biological signals in spite of the variations in epigenetic features, and thereby enabling reliable 3D chromatin structure prediction in diverse biological contexts.

We further evaluated the generalization ability of Chrombus model by TAD as described previously. In all cases, Chrombus models trained on one of the three cell lines showed consistent, significant discrimination power for “between” and “within” TAD interactions in external validation cell lines (Figure 4C).

**Figure 4.**
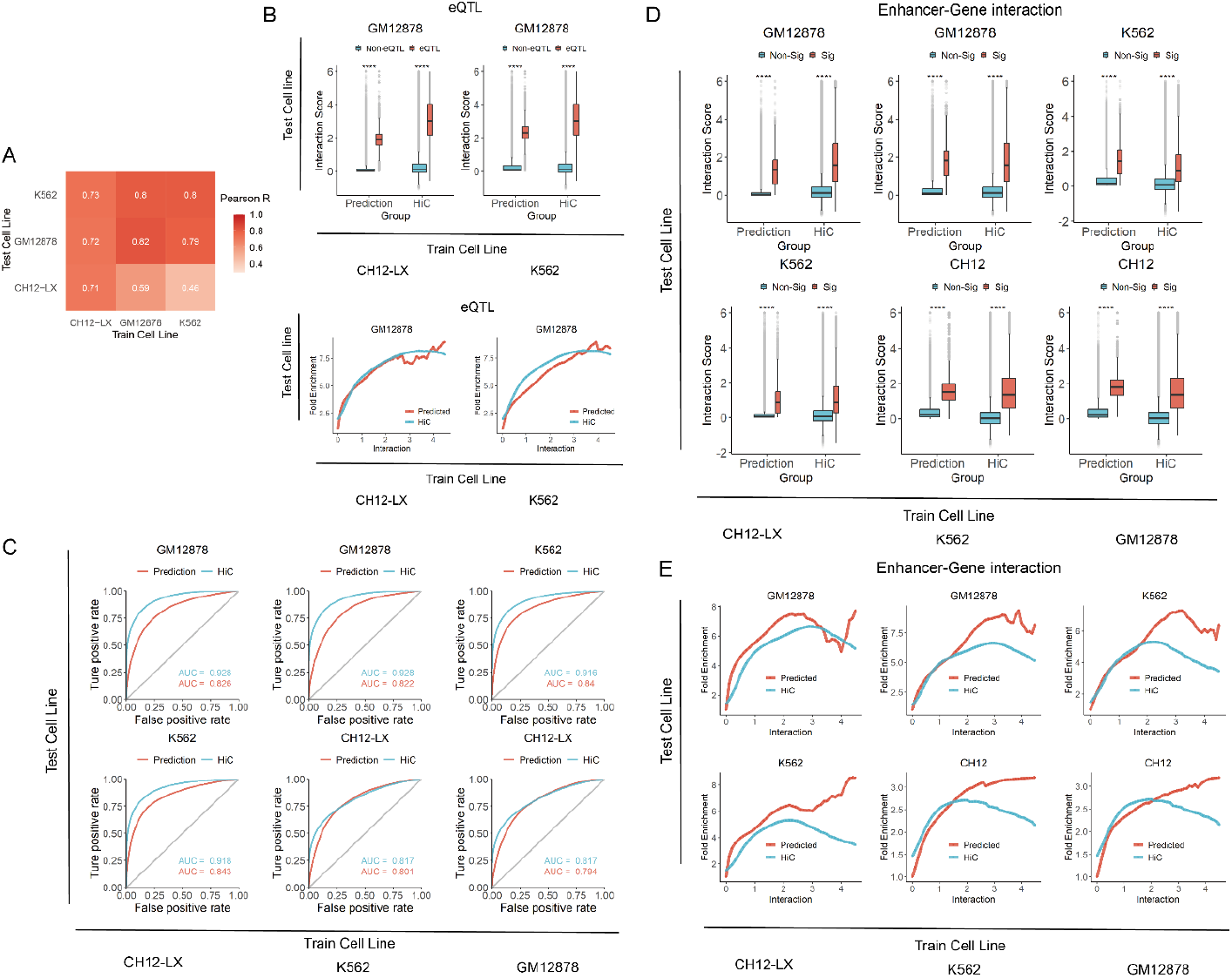
Chrombus is generalizable among different cell linage and species. A: Pearson correlation coefficients between Hi-C scores and predicted scores from 5-fold cross validation in three cell lines, GM12878, K562 and CH12. For each round of validation, the model parameters are trained by one cell line and tested by another. B: Comparison of the interaction scores by Chrombus and Hi-C for segment pairs encompassing eQTL interaction and those do not in GM12878 cell line (upper); fold of enrichment for known enhancer-gene interactions corresponding to different thresholding of the predicted scores (lower). In both cases, the model parameters were train by different cell lines. C: ROC curves for cross-cell-line classification of within- and between-TAD chromatin interactions using predicted scores and Hi-C scores. Again, the model parameters are trained using different cell lines. The TAD loci used as reference are predicted by Arrowhead. D: Comparison of the interaction scores by Chrombus and Hi-C for segment pairs encompassing enhancer-gene interactions with those do not. E: Fold of enrichment for enhancer-gene interactions corresponding to different thresholding of the predicted scores. In all cases the model parameters used for prediction in one cell line were trained by another.

Moreover, Chrombus model trained on one cell line is capable to explain eQTL and enhancer interactions in external cell lines. For instance, when using model trained on GM12878 to predict CTCF-segment pairs in CH12 and K562, we yielded consistently higher predicted scores for CTCF-segments encompassing eQTL interactions and enhancer interactions, mirroring the trends observed in the HiC scores (Figure 4B, D). Again, we noticed that the enrichment of enhancer interactions in the CTCF-segment pairs increases faster with Chrombus prediction scores than with the original HiC scorers, independent of the training set (Figure 4B, E).

These observations imply the persistent enhancement effect of the Chrombus model during model generalization task. Although some variations were observed when extending predictions to mouse cell lines, the model successfully aligns with Hi-C scores and displays promising generalization capabilities, making it a valuable tool in deciphering chromatin interactions across different biological contexts.

### Chrombus is capable of predicting chromatin interactions over one megabase

Currently, only one existing algorithm predict 3D-genome from chromatin features[21]. HiC-Reg uses Random Forest to predict chromatin interactions for 5kb-segmented genomic regions. Here, we compared the predictive performances of HiC-Reg and Chrombus in GM12878 for chromosome 14 and 17.

Within 0.5Mb, HiC-Reg results showed slightly better correlation with Hi-C score than Chrombus did, of which the Pearson’s correlation coefficients mounted to 0.95 and 0.94 from chromosomes 14 and 17, respectively (P < 2.2e-16, Figure 5A, D), while the PCC for Chrombus predictions range from 0.74-0.8 (P < 2.2e-16, Figure 5B, E). As for chromatin interactions between segment pairs of 0.5 to 1Mb apart, the prediction accuracy of HiC-Reg declined significantly, with corresponding PCC of 0.48 and 0.43 (Figure 5A, D). In contrast, Chrombus precition showed a relatively better consistency to Hi-C scores, of which the PCC mounted to 0.58 (chr14) and 0.54 (chr17). Moreover, Chrombus is able to predict long-range chromatin interactions over 1Mb with a stable performance similar to that observed for shorter range, which gives Chrombus the unique advantage of predicting long-range chromatin interactions. The Pearson correlation coefficients ranged 0.51-0.63 (P < 2.2e-16, Figure 5B, E).

**Figure 5.**
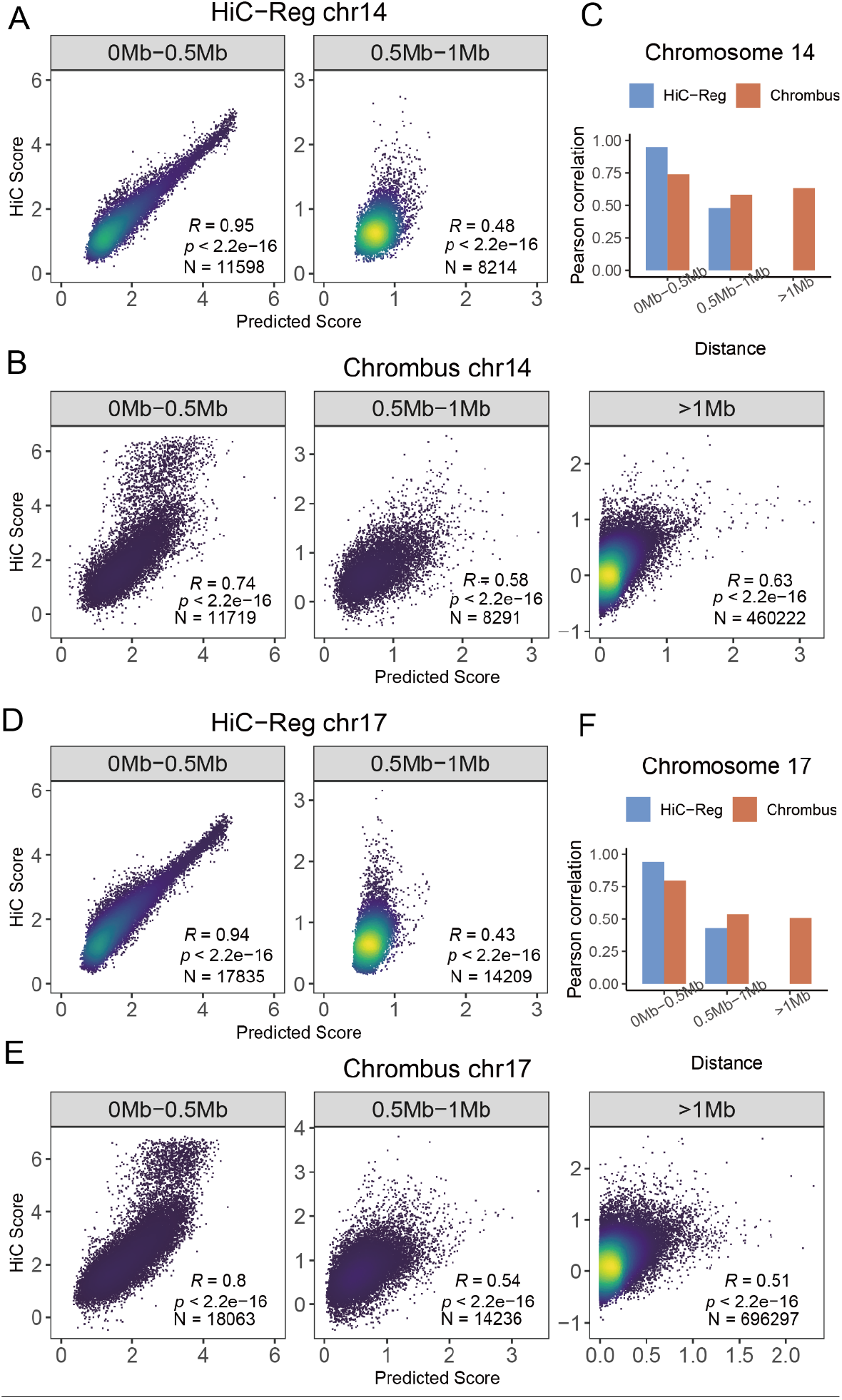
Comparison of the performance of *ab initio* prediction for chromatin interactions by Chrombus and HiC-Reg. A: The correlation between HiC-Reg prediction and Hi-C score for chromatin interactions within 0.5Mb and between 0.5 to 1Mb (chr14). B: The correlation between Chrombus prediction and Hi-C score for chromatin interactions within 0.5Mb, between 0.5 to 1Mb and over 1Mb (chr14). C: Comparison of the Pearson’s correlation coefficients between predicted score and Hi-C score in chromosome 14. The correlation between Hi-C score and predictions by HiC-Reg (D) and Chrombus (E) in chr17 were compared in the same three groups. And the Pearson’s correlation coefficients were also compared (F)

Overall, Chrombus demonstrated comparable performance in predicting short-range interactions while complementing the limitations of similar methods in predicting long-range interactions (Figure 5C, F).

## Conclusion

We described Chrombus-XMBD, a graph convolutional generative model for *ab initio* prediction of 3D-genome. While few methods are developed to resolve 3D-genome based on chromatin features, Chrombus outperforms other methods in two major aspects: first, Chrombus convincingly predicts long-range interactions over 1Mb and second, Chrombus demonstrated exceptional generalizability cross different lineages, suggesting its parameter sets capture highly conserved biological information. Overall, Chrombus-XMBD provides an *in silico* alternative to study *cis-*regulatory mechanisms of gene expression in eukaryote cells.

## Discussion

Resolving the 3D genome offers great advantages to the understanding of the *cis*-regulation of gene expression in eukaryotes. Although many experimental methods have been developed to depict local or genome-wide landscape of chromatin interactions, retrieving the contact-map in the context of heterogeneous tissue or diseases remains challenging. Recent advances in deep learning yielded many successes in solving the dynamics of biomolecules, such as protein 3D-structure, ligand-receptor docking[44, 45]. These methodologies provide an alternative approach of *in silico* prediction of the 3D-conformation of genome based on more accessible features, such as DNA sequences and epigenetic marks. Indeed, several deep-learning methods have demonstrated the potential of using deep learning methods to predict DNA loops [46-49].

The organization of 3D-genome involves complex biological processes. The bonding of nucleotides, recognition and binding of trans-acting proteins such as CTCF and RAD21 (cohesin) to specific DNA motifs are known to be crucial to the formation of DNA-loops[50]. However, the current experimental methods do not support quantifiable measures of the dynamics of DNA-looping which is essential for theoretical statistical modeling. Deep Learning (DL) based methods are data-driven and capable of approximation of complex functions without prior knowledge, which makes it quite suitable for solving 3D genome.

We propose a novel, generative graph convolutional model, Chrombus-XMBD, to predict chromatin interactions using readily available and cost-effective epigenetic data. Our approach interprets chromatin interactions as a graph, which allows for extraction and aggregation of features from interacting DNA segments. Then, Chrombus uses CTCF binding to partition chromatin into unevenly sized DNA fragments, which serve as elementary units of interaction. Chrombus also introduces distance-based regularization on the conventional QKV-attention leverages, which enable sensitive detection of the pattern of chromatin interactions. These characteristics of Chrombus make it stand out of existing methodologies hence offer complementary predictive power to previous models.

Chromatin features are crucial for predicting the 3D genome, but not until recently, few models predict chromatin interactions sole by chromatin features. Chrombus uses six common features: DNA accessibility, CTCF, RAD21, POLR2A, H3K4me3, and H3K27ac, compared with DNA accessibility, CTCF, RAD21, TBP, H3K27me3, H3K9me3, H3K36me3, H4K20me1, H3K79me2, H3K4me1, H3K27ac, H3K9ac, H3K4me2, H3K4me3 by HiC-Reg. Our data suggest the genomic location or vicinity is the most deterministic of chromatin interactions, followed by CTCF binding, H3K27ac, H3K4me3, POLR2A binding and DNA accessibility. Whereas the CTCF motif strand, CTCF motif score and RAD21 (cohesin) show less importance. The effects of the chromatin features are in line with known biology of 3D genome. However, other new chromatin features are yet to be discovered to further improve the prediction.

Of note, Chrombus demonstrated decent predictive power for interactions spanning over 1Mb. As chromatin interaction activity is strongly distance-driven, such long-range interactions are more informative of non-thermal dynamical DNA-looping and implicate specific transcription regulatory events hence more important in biology. In addition, Chrombus are proved to be highly generalizable, which makes it suitable for prediction 3D genome in different cell or tissue types.

Nevertheless, Chrombus-XMBD is limited in for its training data. The availability and quality of the chromatin features are highly variable among databases, platforms and sample types. Owing to its good generalizability, Chrombus can be readily applied to training data of missing features or different noise structure by transfer learning.

Moreover, the experimental technique and the analytical pipeline for Hi-C is not fully matured [51]. The training and validation of Chrombus is subject to unknown biases or noises. Currently, we use TAD inferred from Hi-C, eQTLs and enhancer-gene interactions to validate Chrombus predictions but the most straight forward evidence should base on experimental results such as 3C and 4C.

In summary, we described Chrombus-XMBD, a graph deep learning-based generative model predicts 3D genome *ab intio*. The unique advantages of Chrombus greatly facilitate the exploration of the intricate regulatory mechanisms underlying 3D chromatin structure and the consequential impacts on transcription.

## Reference

1. Zheng, H. and W. Xie, The role of 3D genome organization in development and cell differentiation. Nat Rev Mol Cell Biol, 2019. 20(9): p. 535–550.

2. Kadauke, S. and G.A. Blobel, Chromatin loops in gene regulation. Biochim Biophys Acta, 2009. 1789(1): p. 17–25.

3. Xia, J.H. and G.H. Wei, Enhancer Dysfunction in 3D Genome and Disease. Cells, 2019. 8(10).

4. Zelenka, T., et al., The 3D enhancer network of the developing T cell genome is shaped by SATB1. Nat Commun, 2022. 13(1): p. 6954.

5. Dubois, F., et al., Structural variations in cancer and the 3D genome. Nat Rev Cancer, 2022.22(9): p. 533–546.

6. Tan, L., et al., Changes in genome architecture and transcriptional dynamics progress independently of sensory experience during post-natal brain development. Cell, 2021. 184(3): p. 741–758 e17.

7. Stadhouders, R., G.J. Filion, and T. Graf, Transcription factors and 3D genome conformation in cell-fate decisions. Nature, 2019. 569(7756): p. 345–354.

8. Schoenfelder, S. and P. Fraser, Long-range enhancer-promoter contacts in gene expression control. Nat Rev Genet, 2019. 20(8): p. 437–455.

9. Furlong, E.E.M. and M. Levine, Developmental enhancers and chromosome topology. Science, 2018. 361(6409): p. 1341–1345.

10. Guo, Y., et al., CRISPR Inversion of CTCF Sites Alters Genome Topology and Enhancer/Promoter Function. Cell, 2015. 162(4): p. 900–10.

11. Nichols, M.H. and V.G. Corces, A CTCF Code for 3D Genome Architecture. Cell, 2015.162(4): p. 703–5.

12. Davidson, I.F., et al., Rapid movement and transcriptional re-localization of human cohesin on DNA. EMBO J, 2016. 35(24): p. 2671–2685.

13. Kanke, M., et al., Cohesin acetylation and Wapl-Pds5 oppositely regulate translocation of cohesin along DNA. EMBO J, 2016. 35(24): p. 2686–2698.

14. Ganji, M., et al., Real-time imaging of DNA loop extrusion by condensin. Science, 2018. 360(6384): p. 102–105.

15. Fudenberg, G., et al., Formation of Chromosomal Domains by Loop Extrusion. Cell Rep, 2016. 15(9): p. 2038–49.

16. Tang, Z., et al., CTCF-Mediated Human 3D Genome Architecture Reveals Chromatin Topology for Transcription. Cell, 2015. 163(7): p. 1611–27.

17. Rowley, M.J. and V.G. Corces, Organizational principles of 3D genome architecture. Nat Rev Genet, 2018. 19(12): p. 789–800.

18. Wei, C., et al., CTCF organizes inter-A compartment interactions through RYBP-dependent phase separation. Cell Res, 2022. 32(8): p. 744–760.

19. Fudenberg, G., D.R. Kelley, and K.S. Pollard, Predicting 3D genome folding from DNA sequence with Akita. Nat Methods, 2020. 17(11): p. 1111–1117.

20. Schwessinger, R., et al., DeepC: predicting 3D genome folding using megabase-scale transfer learning. Nat Methods, 2020. 17(11): p. 1118–1124.

21. Zhang, S., et al., In silico prediction of high-resolution Hi-C interaction matrices. Nat Commun, 2019. 10(1): p. 5449.

22. Yang, R., et al., Epiphany: predicting Hi-C contact maps from 1D epigenomic signals. BioRxiv, 2021.

23. Schoenfelder, S., et al., The pluripotent regulatory circuitry connecting promoters to their long-range interacting elements. Genome Res, 2015. 25(4): p. 582–97.

24. Barabasi, A.L. and Z.N. Oltvai, Network biology: understanding the cell’s functional organization. Nat Rev Genet, 2004. 5(2): p. 101–13.

25. The Gene Ontology, C., The Gene Ontology Resource: 20 years and still GOing strong. Nucleic Acids Res, 2019. 47(D1): p. D330–D338.

26. UniProt, C., UniProt: the Universal Protein Knowledgebase in 2023. Nucleic Acids Res, 2023. 51(D1): p. D523–D531.

27. Gao, Z., et al., Hierarchical graph learning for protein-protein interaction. Nat Commun, 2023. 14(1): p. 1093.

28. Munoz, E., V. Novacek, and P.Y. Vandenbussche, Facilitating prediction of adverse drug reactions by using knowledge graphs and multi-label learning models. Brief Bioinform, 2019. 20(1): p. 190–202.

29. Consortium, E.P., et al., Expanded encyclopaedias of DNA elements in the human and mouse genomes. Nature, 2020. 583(7818): p. 699–710.

30. Dekker, J., et al., The 4D nucleome project. Nature, 2017. 549(7671): p. 219–226.

31. Rao, S.S., et al., A 3D map of the human genome at kilobase resolution reveals principles of chromatin looping. Cell, 2014. 159(7): p. 1665–80.

32. Grant, C.E., T.L. Bailey, and W.S. Noble, FIMO: scanning for occurrences of a given motif. Bioinformatics, 2011. 27(7): p. 1017–8.

33. Thomas N. Kipf, M.W., Variational Graph Auto-Encoders. 1611.07308, 2016.

34. Ashish Vaswani, N.S., Niki Parmar, Jakob Uszkoreit, Llion Jones, Aidan N. Gomez, Lukasz Kaiser, Illia Polosukhin, Attention Is All You Need. 1706.03762, 2017.

35. Wolff, J., et al., Galaxy HiCExplorer 3: a web server for reproducible Hi-C, capture Hi-C and single-cell Hi-C data analysis, quality control and visualization. Nucleic Acids Res, 2020. 48(W1): p. W177–W184.

36. Consortium, G.T., The GTEx Consortium atlas of genetic regulatory effects across human tissues. Science, 2020. 369(6509): p. 1318–1330.

37. Fishilevich, S., et al., GeneHancer: genome-wide integration of enhancers and target genes in GeneCards. Database (Oxford), 2017. 2017.

38. van Schoonhoven, A., et al., 3D genome organization during lymphocyte development and activation. Brief Funct Genomics, 2020. 19(2): p. 71–82.

39. Qiu, Y., et al., 3D genome organization and epigenetic regulation in autoimmune diseases. Front Immunol, 2023. 14: p. 1196123.

40. Rodriguez-Carballo, E., et al., The HoxD cluster is a dynamic and resilient TAD boundary controlling the segregation of antagonistic regulatory landscapes. Genes Dev, 2017. 31(22): p. 2264–2281.

41. Xia, Y., et al., Capturing 3D Chromatin Maps of Human Primary Monocytes: Insights From High-Resolution Hi-C. Front Immunol, 2022. 13: p. 837336.

42. Luo, H., et al., HOTTIP lncRNA Promotes Hematopoietic Stem Cell Self-Renewal Leading to AML-like Disease in Mice. Cancer Cell, 2019. 36(6): p. 645–659 e8.

43. Kang, J., et al., Multiple CTCF sites cooperate with each other to maintain a TAD for enhancer-promoter interaction in the beta-globin locus. FASEB J, 2021. 35(8): p. e21768.

44. Jumper, J., et al., Highly accurate protein structure prediction with AlphaFold. Nature, 2021. 596(7873): p. 583–589.

45. Peng, L., et al., Deciphering ligand-receptor-mediated intercellular communication based on ensemble deep learning and the joint scoring strategy from single-cell transcriptomic data. Comput Biol Med, 2023. 163: p. 107137.

46. Lv, H., et al., A sequence-based deep learning approach to predict CTCF-mediated chromatin loop. Brief Bioinform, 2021. 22(5).

47. Trieu, T., A. Martinez-Fundichely, and E. Khurana, DeepMILO: a deep learning approach to predict the impact of non-coding sequence variants on 3D chromatin structure. Genome Biol, 2020. 21(1): p. 79.

48. Dao, F.Y., et al., DeepYY1: a deep learning approach to identify YY1-mediated chromatin loops. Brief Bioinform, 2021. 22(4).

49. Wang, S., et al., DLoopCaller: A deep learning approach for predicting genome-wide chromatin loops by integrating accessible chromatin landscapes. PLoS Comput Biol, 2022. 18(10): p. e1010572.

50. Sanborn, A.L., et al., Chromatin extrusion explains key features of loop and domain formation in wild-type and engineered genomes. Proc Natl Acad Sci U S A, 2015. 112(47): p. E6456–65.

51. Sefer, E., A comparison of topologically associating domain callers over mammals at high resolution. BMC Bioinformatics, 2022. 23(1): p. 127.

