## Supplemental Table1-2, Supplemental Figure 1-5 Table1-2 for "Chrombus-XMBD: A Graph Generative Model Predicting 3D-Genome, *ab initio* from Chromatin Features"

Supplementary Table 1 Data source

| Data Type | Mark | File format | GM12878 | H1ESC | K562 |
|---|---|---|---|---|---|
| CTCF | Domain Boundary | bed-narrowPeak fold-change-over-control | ENCFF473RXY ENCFF886KRA | ENCFF093VEE ENCFF093CSL | ENCFF738TKN ENCFF933ZLL |
| RAD21 | Cohesin | bed-narrowPeak | ENCFF001VFE | ENCFF348QBX | ENCFF002CXU |
| POLR2A | RNA polymerase II mark | fold-change-over-control | ENCFF368HBX | ENCFF339CZB | ENCFF647MSS |
| H3K4me3 | Activating mark | fold-change-over-control | ENCFF818GNV | ENCFF818GNV | ENCFF291SWG |
| H3K27ac | Enhancer mark | fold-change-over-control | ENCFF180LKW | ENCFF809EWV | ENCFF010PHG |
| DNA accessibility | Open chromatin signal | read-depth normalized signal | ENCFF901GZH | ENCFF935TUU | ENCFF352SET |
| Chromatin interaction | Contact counts | Contact matrix and fastq | GSE63525 | 4DNFID162B9J | GSE63525 |

Supplementary Table 2 Preprocessing of 14-dimensional node features

| **Raw data** | **Preprocessing** | **Feature** | **Raw value** | **Normalization** | **Processed value** |
|---|---|---|---|---|---|
| CTCF ChIP-Seq | fimo | Strand of CTCF motif (left) | Discrete value (“+” / “-”) from fimo | Convert “+” to 1 and “-” to 0 | 1 / 0 |
| | | Scores of CTCF motif (left) | Continuous numeric value (> 0) from fimo | Min-max | Range (0, 1) |
| | | Strand of CTCF motif (right) | Discrete value (“+” / “-”) from fimo | Convert “+” to 1 and “-” to 0 | 1 / 0 |
| | | Scores of CTCF motif (right) | Continuous numeric value (> 0) from fimo | Min-max | Range (0, 1) |
| CTCF ChIP-Seq | fold-change-over-control | CTCF binding peak signal (left) | Continuous numeric value (> 0) | Min-max | Range (0, 1) |

| | | | | | |
|---|---|---|---|---|---|
| | | CTCF binding peak signal (right) | Continuous numeric value (> 0) | Min-max | Range (0, 1) |
| Histone ChIP-Seq | | H3K27ac signal | Continuous numeric value (> 0) | Min-max | Range (0, 1) |
| | | H3K4me3 signal | Continuous numeric value (> 0) | Min-max | Range (0, 1) |
| TF ChIP-Seq | | POLR2A signal | Continuous numeric value (> 0) | Min-max | Range (0, 1) |
| DNase-Seq | | DNase I signal | Continuous numeric value (> 0) | Min-max | Range (0, 1) |
| TF ChIP-Seq | bed-narrowPeak | RAD21 binding (left) | Boolean | Convert “True” to 1 and “False” to 0 | 1 / 0 |
| | | RAD21 binding (right) | Boolean | Convert “True” to 1 and “False” to 0 | 1 / 0 |
| Hg19 genome building | | Start position of segment | Integer (> 0) | Position / length of chromosome | Range (0, 1) |
| | | End position of segment | Integer (> 0) | Position / length of chromosome | Range (0, 1) |

Supplementary Figure 1 Choose methods for normalizing contact counts

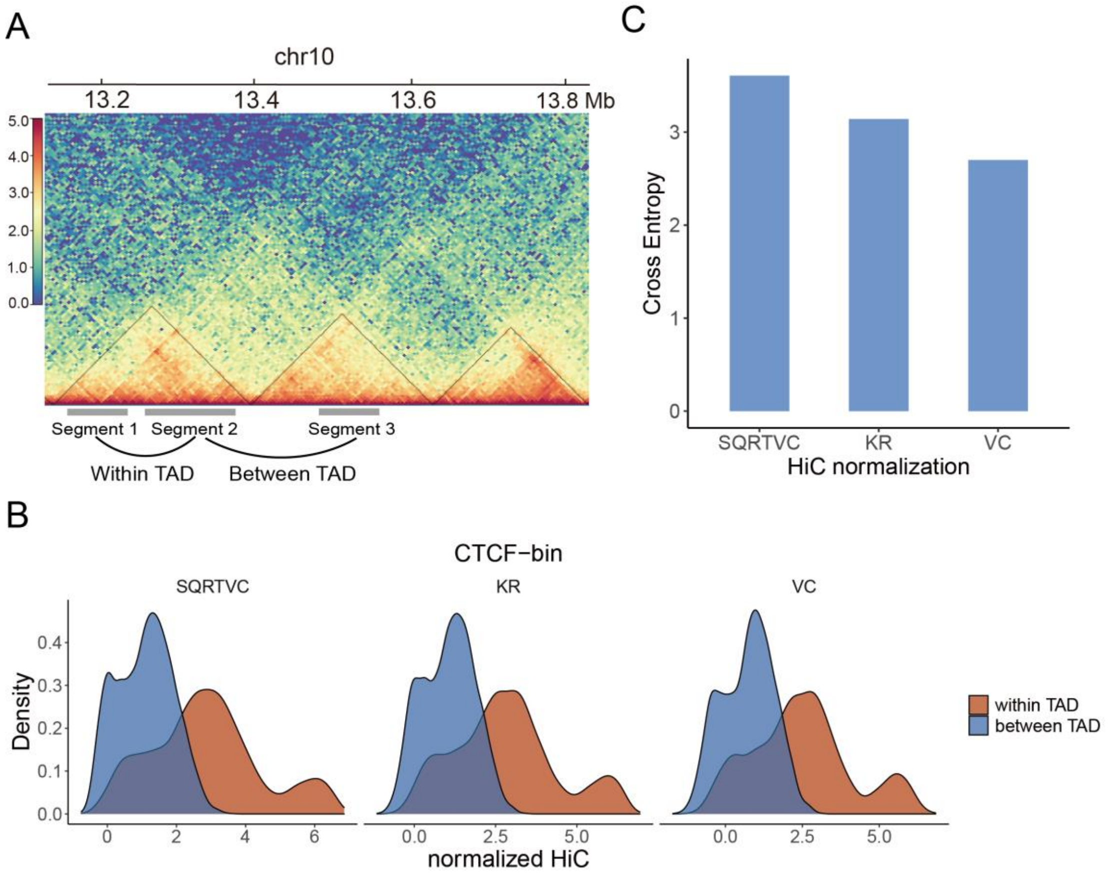


Supplementary Figure 2 Distribution of input features

C

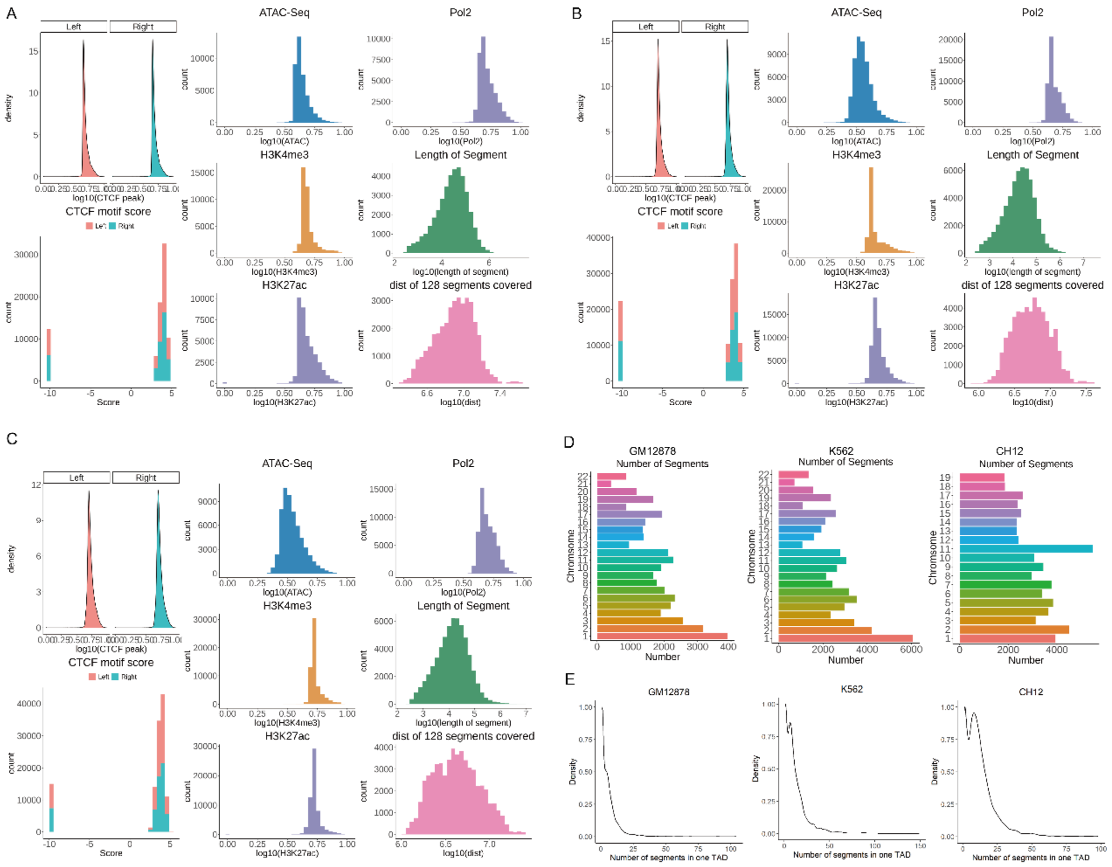


Supplementary Figure 3 Training loss of GM12878 model (A) and correlation of predicted score and HiC score (B).

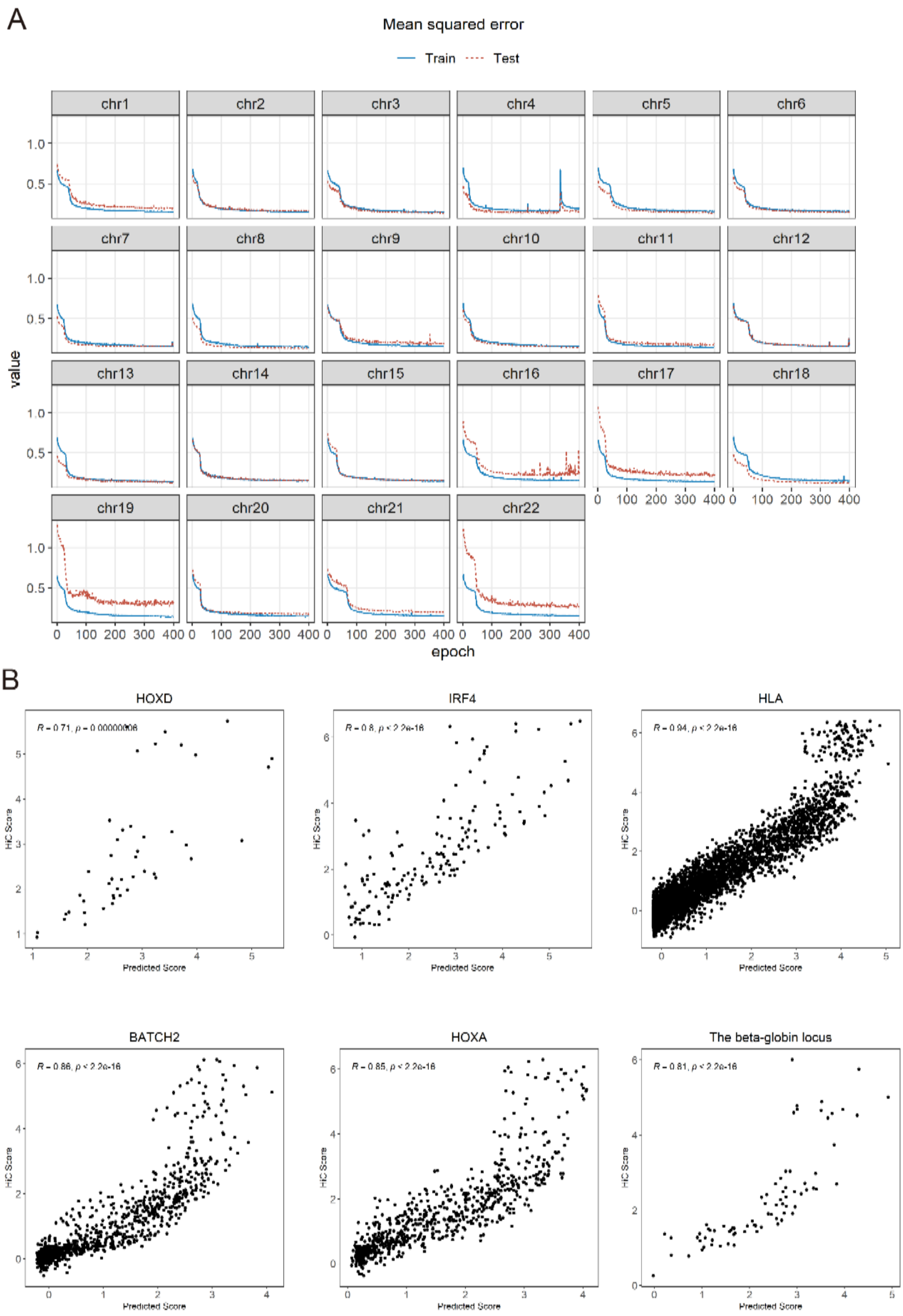


Supplementary Figure 4 K-fold cross validation model (K=5)

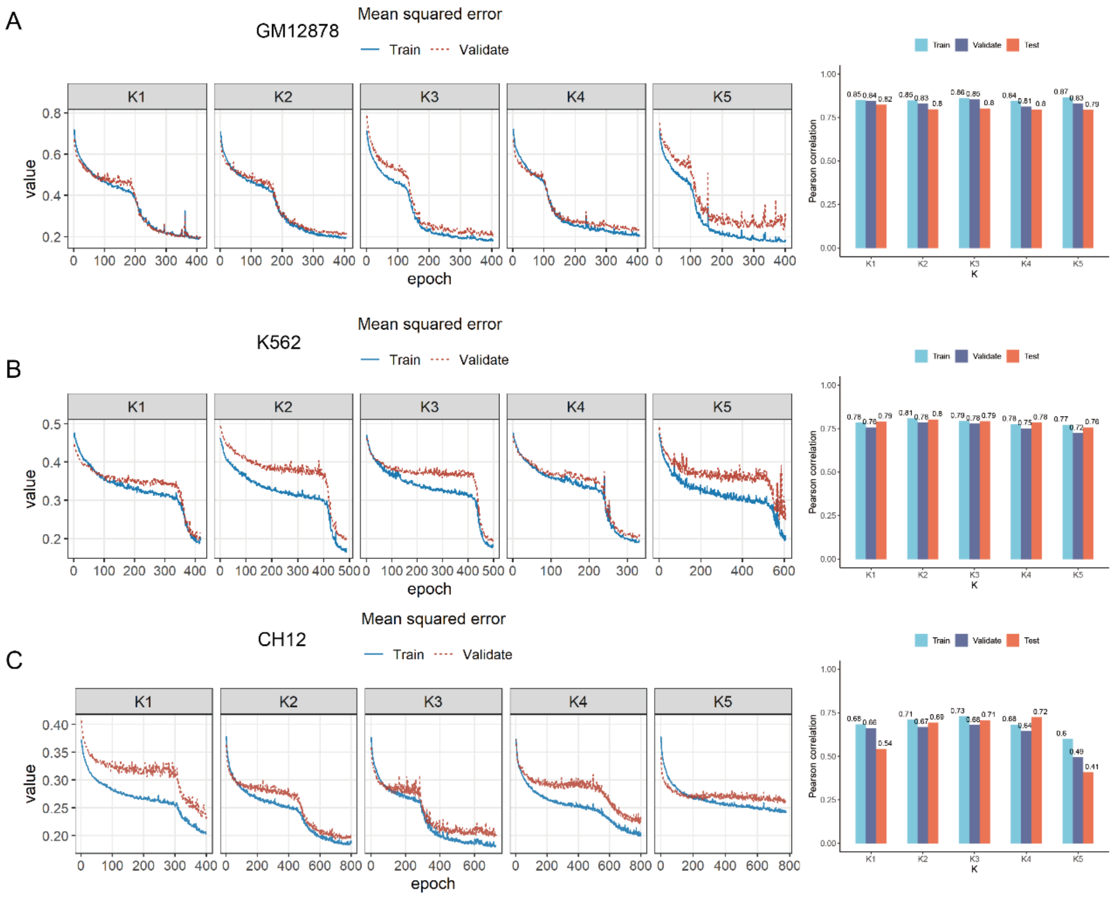


Supplementary Figure 5 Feature importance (A) and learned node representation from Chrombus

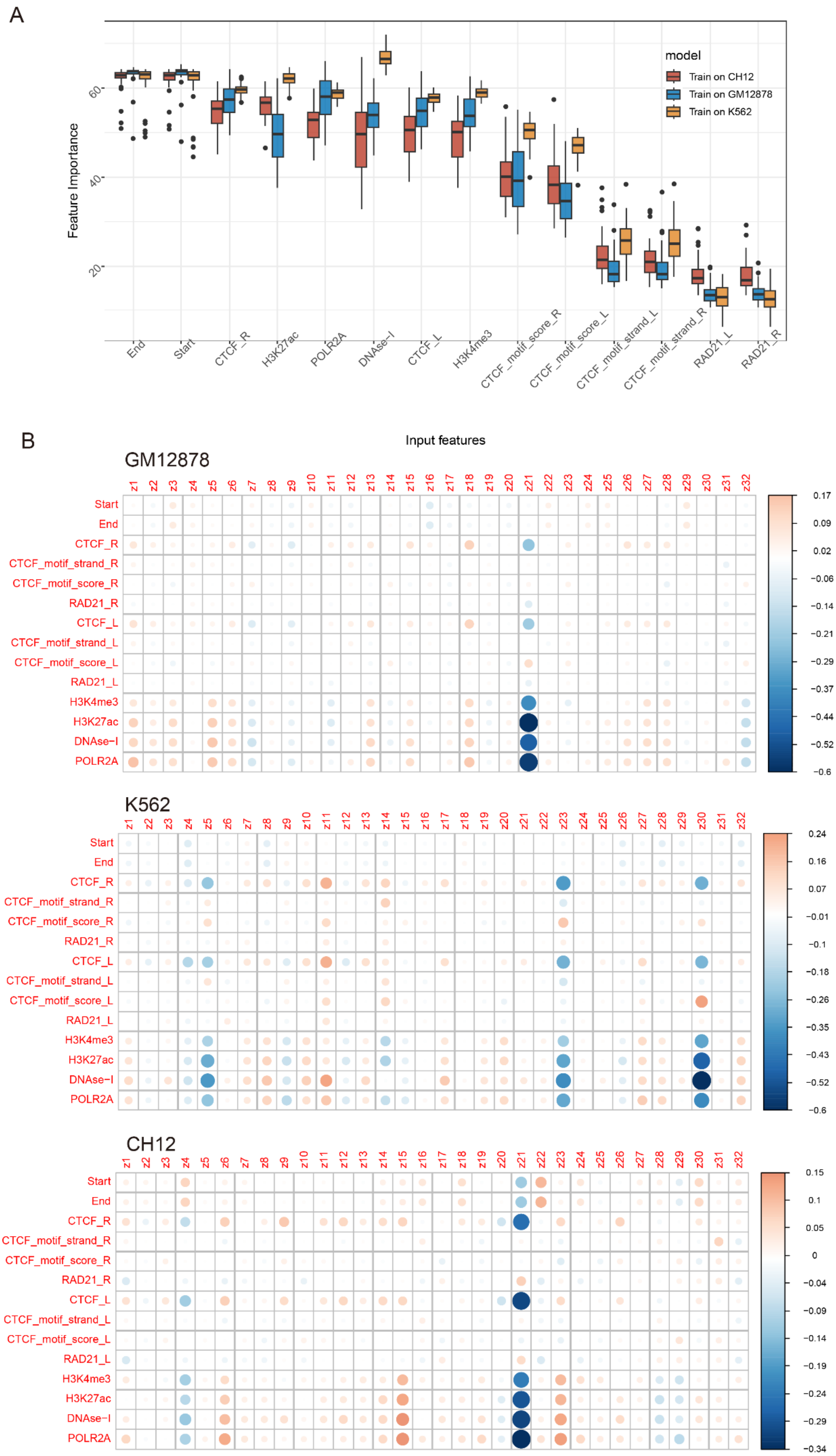

A
Feature Importance
20
40
60
End
Start
CTCF_R
H3K27ac
POLR2A
DNAse-I
CTCF_L
H3K4me3
CTCF_motif_score_R
CTCF_motif_score_L
CTCF_motif_strand_L
CTCF_motif_strand_R
RAD21_L
RAD21_R
model
Train on CH12
Train on GM12878
Train on K562
B
Input features
GM12878
K562
CH12
z1 z2 z3 z4 z5 z6 z7 z8 z9 z10 z11 z12 z13 z14 z15 z16 z17 z18 z19 z20 z21 z22 z23 z24 z25 z26 z27 z28 z29 z30 z31 z32
Start
End
CTCF_R
CTCF_motif_strand_R
CTCF_motif_score_R
RAD21_R
CTCF_L
CTCF_motif_strand_L
CTCF_motif_score_L
RAD21_L
H3K4me3
H3K27ac
DNAse-I
POLR2A
0.17 0.09 0.02 −0.06 −0.14 −0.21 −0.29 −0.37 −0.44 −0.52 −0.6
0.24 0.16 0.07 −0.01 −0.1 −0.18 −0.26 −0.35 −0.43 −0.52 −0.6
0.15 0.1 0.05 0 −0.04 −0.09 −0.14 −0.19 −0.24 −0.29 −0.34
